# Psilocybin-induced modulation of visual salience processing

**DOI:** 10.1101/2025.05.10.652897

**Authors:** Stephanie Muller, Federico Cavanna, Laura Alethia de la Fuente, Nicolás Bruno, Tomás Ariel D’Amelio, Carla Pallavicini, Enzo Tagliazucchi

## Abstract

Psychedelic compounds significantly reshape conscious perception, yet the implications of these alterations for complex visual-guided behaviors remain poorly understood. We investigated how psilocybin modulates visual salience processing during natural scene perception. Twenty-three participants completed eye-tracking tasks under self-blinded low and high doses of psilocybin, in a naturalistic design with experimental conditions unknown to participants and researchers. Subjects viewed natural scenes while their gaze patterns were recorded and analyzed in relation to normative computational saliency maps generated using a deep learning model of visual attention. Results revealed increased fixation on salient image regions and reduced inter-fixation distance under the high-dose condition, suggesting heightened sensitivity to visual salience and more localized gaze behavior. The Shannon entropy of fixations on high-saliency regions indicated a more exploratory and less predictable visual scanning of the images. Complementary EEG recordings showed broadband spectral power reductions and increased Lempel-Ziv complexity, with delta power negatively correlating with salience metrics. These findings suggest psilocybin enhances bottom-up attentional control while weakening top-down modulation, consistent with theoretical models positing facilitated bottom-up information flow under the acute effect of psychedelics.

## Introduction

Serotonergic psychedelics are notable for their ability to induce profound alterations in sensory perception (Nichols, 2016; van Elk & Yaden, 2022). In the visual domain, psychedelic tryptamines such as psilocybin and dimethyltryptamine induce low-level alterations, including changes in the color and texture of objects and the appearance of geometric distortions (Preller & Vollenweider, 2016; Kometer & Vollenweider, 2018; Aday et al., 2021). Despite these widespread effects, there are relatively few studies investigating how visual alterations impact complex behaviors that depend on the integration of multiple cognitive functions (Bălăeţ, 2022). One particularly underexplored area is the impact of psychedelics on attentional processes involved in integrating visual information for object recognition in natural scenes.

Humans navigate their environment by continuously sampling visual information through a sophisticated pattern of eye movements, including saccades, fixations, and microsaccades, collectively enabling the dynamic exploration and processing of visual scenes (Findlay & Gilchrist, 2003). The neural and temporal costs associated with the serial nature of fixations require mechanisms for selecting certain aspects of the image over others (Gottlieb et al., 2014). Bottom-up prioritization is driven by visual salience and depends on low-level image features such as specific colors, shapes, contrast, and motion. Fixation patterns are also modulated by top-down factors including task-demands, higher-order cognitive processes, and individual variations. Bottom-up and top-down influences interact dynamically to guide visual attention, balancing sensory-driven responses with goal-directed control of gaze (Fecteau & Munoz, 2006; Tatler et al., 2011).

Psychedelic substances can influence this process at multiple instances in parallel, with an overall uncertain effect. Perceptual distortions may affect the identification of low-level information in the visual scene, interfering with the formation of saliency maps and basic feature detection. These alterations include changes in contrast sensitivity, color perception, and edge detection, which could fundamentally alter how visual information is processed in early visual areas (Kometer & Vollenweider, 2018). Psychedelics have also been shown to broaden visual perceptual bandwidth, as evidenced by increased prepulse inhibition (Quednow et al., 2012; Vollenweider et al., 2007). Impaired evaluation of sensory signal accuracies can result in a more disorganized or entropic exploration of the visual scene (Carhart-Harris et al., 2014). Additionally, changes in attention allocation and cognitive control mechanisms might contribute to more erratic scanning patterns and modified fixation strategies (Bălăeţ, 2022; Carter et al., 2005; Quednow et al., 2012)

The present study investigated the effects of psilocybin on visual salience processing during natural scene perception. Following a randomized self-blinded protocol conducted in ecologically valid settings, participants completed two dosing sessions—one low-dose and one high-dose of psilocybin mushrooms on separate days and were subsequently presented with natural scenes for brief time intervals. The sequence of gaze fixations was recorded using an eye-tracking device, informing how the subjects prioritized specific regions of the images. In combination with self-reported psychometric data, resting-state electroencephalography (EEG) was recorded to obtain spectral markers of subjective effect intensity, and to investigate the neural correlates of psilocybin-induced modulation of visual salience processing.

We examined four primary hypotheses regarding the impact of psilocybin on visual perception and attention. First, we hypothesized changes in salience associated with gaze fixations under psilocybin, in relation to a normative model of spatial salience calibrated with data from individuals in a normal state of consciousness. Second, we hypothesized increased entropy in visual exploration patterns across high salience regions, quantified by the entropy of fixations among the individual regions of highest salience in the image. Third, we anticipated a broadband reduction in EEG spectral power, hypothesized to correlate with the behavioral measures outlined in our initial predictions. Lastly, based on our previous findings regarding aesthetic perception (Muller et al., 2023), we predicted specific alterations in general visual exploration strategies, particularly a more localized exploration of the visual scene.

## Methods

### Participants

Twenty-three participants (four females, 31 ± 4 years, 72 ± 15 kg [mean ± STD]) were recruited through word of mouth and social media advertising. To participate, individuals were instructed to contact the provided number via WhatsApp and subsequently received a phone call from the researchers. During the call, the researchers provided a brief explanation of the details and purpose of the experiment, as well as the inclusion and exclusion criteria. Participants were then given a full written explanation of the study and a copy of the informed consent form. To determine eligibility, all participants underwent a psychiatric interview to screen for exclusion criteria, which are detailed in the supplementary material and summarized below. After this screening, participants and researchers agreed on a date for the start of the experiment. Participants reported having 15 ± 13 prior experiences with serotonergic psychedelics, of which 3.1 ± 2.4 were considered challenging [mean ± STD]. All participants had normal or corrected-to-normal vision.

This study was conducted in accordance with the Declaration of Helsinki and approved by the Research Ethics Committee at the Universidad Abierta Interamericana (Buenos Aires, Argentina), protocol number 0-1068. All participants gave written informed consent and received no financial compensation for their participation in the experiment. All data was collected from the participants in natural settings, i.e. those chosen by the participants without intervention from the researchers. The researchers did not provide nor administered psilocybin to the participants, nor instructed them in any way concerning drug use.

### Inclusion and exclusion criteria

Participants were required to have at least two prior experiences with a dose equal to or exceeding 3 g of dried psilocybin mushrooms. To participate in this research protocol, subjects volunteered to partake in a series of tests under the effects of psilocybin mushrooms and in the presence of four members of the research team. Subjects who consumed serotonergic psychedelics during the 15 days prior to the dosing day were not included in the study. The same applied for all psychoactive substances (including alcohol, caffeine and tobacco) for a period of 24 hours prior to the dosing day. To participate in the experiment, subjects declared their willingness to abstain from using psychedelics between measurement sessions.

Subjects who fulfilled DSM-5 criteria for the following disorders were excluded from the experiment: schizophrenia or other psychotic disorders, type 1 or 2 bipolar disorder (including first- and second-degree relatives), personality disorders, dissociative identity disorder, post-traumatic stress disorder, substance abuse or dependence in the past 5 years, depressive disorders, recurrent depressive episodes, obsessive-compulsive disorder, generalized anxiety disorder, dysthymia, panic disorder, bulimia or anorexia, as well as subjects with a history of neurological disorders. Pregnant women and subjects under psychiatric medication of any kind were excluded. Subjects exhibiting potential dysfunctional states as measured by the Depression Anxiety Stress Scale (DASS) (with scores > 4 for depression, > 3 for anxiety and > 7 for stress) were subsequently reviewed by the clinical interviewer for confirmation.

### Experimental design and setting

Experimental conditions (high vs. low dose) were a priori unknown to participants and researchers as subjects implemented a self-blinding procedure, following the design introduced by and colleagues (2021). The experiment was divided into two parts, one corresponding to the dosing condition (3 g of ground and homogenized dried psilocybin mushrooms in gel capsules, provided by the participants) and one corresponding to an active control condition (0.5 g of ground and homogenized dried psilocybin mushrooms mixed with edible mushrooms to match the weight of the gel capsules between conditions). Conditions were separated by an interval of one month to attenuate potential tolerance effects. During interviews conducted after completion of the experiment, we confirmed that participants implemented the self-blinding assisted by a third party of their choice. In all cases, experiments took place in the comfortable setting of a house and in the presence of the participant and the team of researchers. During the high-dose session, two participants opted to discontinue the eye-tracking task. Consequently, they were excluded from subsequent analyses of eye-tracking data, although no adverse effects were reported.

### Self-reported scales and questionnaires

Two days before the dosing day for each condition, participants completed a set of self-reported questionnaires designed to assess various psychological traits. These included the State-Trait Anxiety Inventory (STAI-Trait) (Spielberger, 1983), the Big Five Inventory (BFI) (John et al., 1991), the Tellegen Absorption Scale (TAS) (Tellegen & Atkinson, 1974), and the Short Suggestibility Scale (SSS) (Kotov, 2004). Additional self-report scales were administered both before dosing and immediately after the acute effects, including the State-Trait Anxiety Inventory (STAI-State) (Spielberger, 1983) the Positive and Negative Affect Schedule (PANAS) (Sandin, 1999), and the Psychological Well-being Scale (BIEPS) (Luna et al., 2020). Before dosing, participants also completed assessments on expectations (EXP) and contextual factors related to prior psychedelic experiences (PRE) (Haijen et al., 2018). Following the acute effects, participants completed the Mystical Experience Scale (MEQ) (MacLean et al., 2011) and the Altered States of Consciousness Scale (5D-ASC) (Studerus et al., 2010). In addition, participants rated subjective drug effects using a visual analogue scale (VAS) at four time points during the dosing day, beginning one hour after ingestion and continuing at hourly intervals until the effects subsided. A comprehensive description of all questionnaires used in the study is provided in the supplementary material.

### Stimuli

A total of 100 images were randomly subsampled from the OSIE dataset (Xu et al., 2014), which originally comprised 700 images. The original OSIE dataset was deliberately designed with the objective of mitigating the biases that are commonly observed in existing image collections. In contrast to traditional datasets, which often feature a single dominant object in a central position, the images in OSIE are characterised by the presence of multiple dominant objects distributed across the scene. The images are diverse, spanning a wide range of object categories with strong semantic relevance. This design supports detailed analyses of attentional dynamics by incorporating both pixel-level details and object- and semantic-level attributes. The selected images had a resolution of 800 × 600 pixels and were displayed against a uniform grey background to ensure consistent viewing conditions.

### Eye tracking

The measurements described below were conducted as part of a broader study examining the effects of psychedelics on perception, creativity, language, and music production. Findings related to these domains will be presented in future reports. Prior to the task described in this study, participants completed another eye-tracking task involving free exploration of artworks from various periods and styles, with the results of this experiment detailed in a previous publication (Muller et al., 2023)

For the eye-tracking tasks, participants were seated 60 cm from a centrally positioned monitor, with their heads stabilized using a custom-made chin rest to minimize movement. The eye tracker was positioned between the participant and the screen. The display measured 31 × 17.5 cm and had a resolution of 1920 × 1080 pixels. During the presentation of visual stimuli, gaze coordinates were recorded in both spatial and temporal dimensions using the Gazepoint GP3 HD portable eye tracker, which operates at a temporal resolution of 150 Hz and achieves a visual angle accuracy of 0.5–1.0 degrees. Stimulus presentation and eye-tracking control were managed by a custom Python script developed using the open-source PsychoPy library (https://www.psychopy.org/).

The task reported in this study began 90 minutes after participants consumed capsules containing either a low or high dose of psilocybin. An overview of the experimental procedure is provided in Figure 1. Each trial (T) commenced with a fixation cross displayed for one second, followed by a stimulus presentation lasting three seconds, during which participants were free to observe the image. The same stimuli were used across all experimental conditions, with their order randomized for each participant and condition. A standard five-point calibration was conducted after every twentieth block to ensure that the eye tracking detected fixations accurately throughout the duration of the experiment.

**Figure 1.**
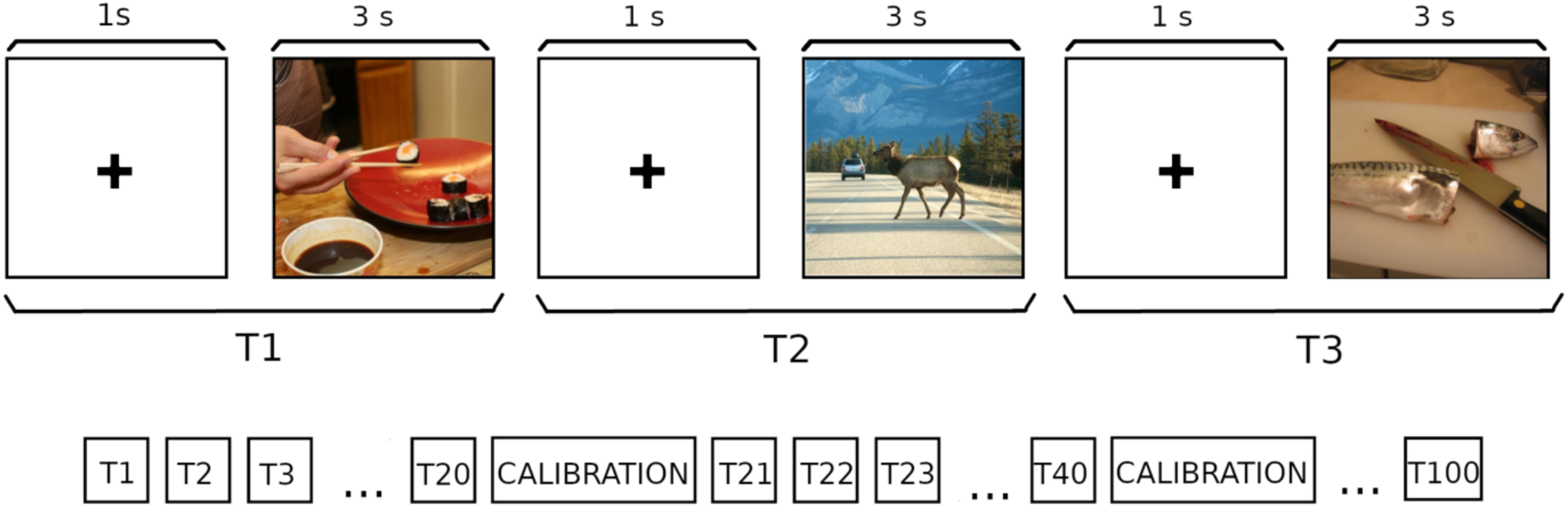
Overview of the eye-tracking task. Each trial consisted of an initial calibration phase, followed by the first block (T1). Each block included the following steps: a fixation cross displayed for 1 second, followed by a 3-second stimulus presentation. During this time, participants freely explored the image while their gaze was recorded by the eye tracker. This procedure was repeated for all 100 images, with intermediate eye tracker calibrations every 20 blocks.

### Statistical fixation metrics

Due to insufficient data quality, i.e. more than 30% of fixations either missing or recorded outside the screen for over 30% of the stimuli (suggestive of lack of engagement with the task), data from six participants was excluded from the present analysis. Furthermore, for each participant and condition, only images with less than 30% of fixations outside the boundaries were analyzed. All fixations beyond the image boundaries were excluded from further analyses. No significant differences were found in the number of excluded fixations across experimental conditions.

A velocity-based algorithm (Engbert & Kliegl, 2003) was employed to automatically identify fixations with minimum durations of 150 ms following the recording of gaze position coordinates (Manor & Gordon, 2003). The mean horizontal (*x_i_*) and vertical (*y_i_*) positions of each fixation, along with the time of the i-th fixation (t*_i_*) were then determined, with positions expressed in pixels and time in seconds. For each image, participant, and condition, the following metrics were calculated: the total number of fixations within the image (N), the mean distance between fixations (ds), and the mean time between fixations (dt). The following formulas were applied to compute these measures:

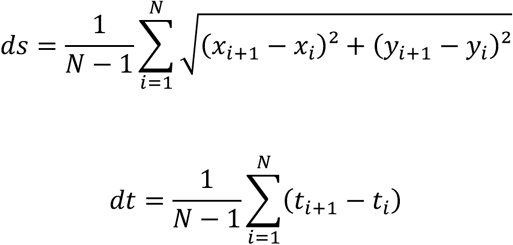

These measures were subsequently averaged across all images, yielding a mean value for each participant under each dosing condition.

### Saliency

To analyze the visual saliency, we first converted each image into a saliency map using DeepGaze II, a computational model of visual attention based on deep neural networks trained to predict human fixations on natural images. The model was pretrained on the SALICON dataset, which contains 10,000 diverse images annotated with mouse-tracking data, and subsequently fine-tuned on the MIT1003 dataset, composed of 1003 natural scene photographs viewed by human participants under eye-tracking for 3 seconds. Validation was conducted via 10-fold cross-validation on the MIT1003 dataset to ensure the model generalizes well to unseen images within the same domain (Linardos et al., 2021). The generated saliency maps reflect the likelihood of a region capturing visual attention. Each recorded fixation was then mapped to its corresponding saliency value from the respective image’s saliency map. The average saliency for each trial was calculated, and a final saliency value for each participant and experimental condition was obtained by averaging across all images. To examine how participants’ focus on salient regions evolved over time, we divided the 3-second interval into five parts and calculated the average saliency of the fixations within each subinterval.

### Shannon Entropy in High Salience Regions

To quantify the predictability of fixation patterns in high saliency regions, we computed the Shannon entropy of fixation distributions within high-saliency areas. The saliency map was first binarized using a threshold, which was systematically varied in subsequent analysis to assess the robustness of the results across different saliency levels. Connected clusters of the binary map were identified and treated as individual regions with high visual salience (see figure 4a). Each fixation was assigned to a high salience cluster based on its spatial location. The Shannon entropy was then calculated for the resulting distribution of fixations across clusters, using the formula:

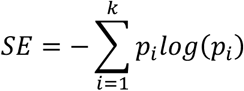

where *p_i_* represents the proportion of fixations within the i-th cluster relative to the total number of fixations within the clusters and *k* is the total number of clusters. Thus, higher entropy values indicate a more uniform distribution of fixations across the high salience clusters of the image.

This analysis only included images with more than one cluster to ensure the entropy calculation accurately reflected variability in fixation patterns. The analysis was performed across a range of thresholds to ensure the results were not biased by the chosen value. The lower bound of the explored range was defined as the threshold at which at least 70% of the images contained more than one cluster, while the upper bound corresponded to the point where the average number of clusters started to decline, indicating a loss of information about salient regions. For each threshold, entropy was compared between dosing conditions.

### EEG acquisition, preprocessing and analysis

Resting state EEG data was recorded using a 24-channel mobile system (mBrainTrain LLC, Belgrade, Serbia; http://www.mbraintrain.com/) paired with an elastic electrode cap (EASYCAP GmbH, Inning, Germany; www.easycap.de). Twenty-four Ag/AgCl electrodes were positioned according to the standard 10–20 system at the following locations: Fp1, Fp2, Fz, F7, F8, FC1, FC2, Cz, C3, C4, T7, T8, CPz, CP1, CP2, CP5, CP6, TP9, TP10, Pz, P3, P4, O1, and O2. Reference and ground electrodes were placed at FCz and AFz, respectively. The wireless EEG DC amplifier (weight: 60 g; dimensions: 82 × 51 × 12 mm; resolution: 24-bit; sampling rate: 500 Hz; passband: 0–250 Hz) was securely attached to the back of the electrode cap, between electrodes O1 and O2. EEG signals were digitized and transmitted via Bluetooth to a notebook operated by the experimenter, who was positioned behind the participant. Participants were instructed to undertake five-minute recordings with their eyes closed, both 15 minutes prior to and two hours after the administration of the dose. During these recordings, the subjects were asked to keep their eyes fully closed, relax all muscles, and avoid any physical tension.

EEG data was preprocessed using the Python MNE library (Gramfort, 2013). A band-pass filter with a frequency range of 0.5–90 Hz was applied to the data to remove low-frequency drift and high-frequency noise. Additionally, a notch filter was employed to eliminate power line noise at 50 Hz and its harmonics. The filtered data were then segmented into fixed-length epochs of two seconds for subsequent analysis. Automatic epoch rejection was conducted using MNE’s autoreject algorithm (Jas et al., 2017), which identified and excluded epochs characterised by excessive noise. A manual inspection was conducted to ensure the quality of the data. Independent component analysis (ICA) was then applied to the cleaned epochs to identify components that were associated with artefacts such as eye blinks and muscle activity. This was subsequently corroborated by visual inspection. The remaining bad channels were interpolated, and the data was re-referenced to the average of all channels. According to the previously defined criteria, three subjects were excluded from the subsequent EEG analysis due to an excessive number of rejected epochs and/or channels in at least one condition. This resulted in 20 subjects for subsequent analysis, with a mean of 119 ± 22 epochs.

The power spectral density (PSD) was computed for each participant and experimental condition using the *MNE* implementation, applying the multitaper method. The minimum and maximum frequencies were set to 1 Hz and 40 Hz, respectively. PSDs were averaged across epochs and channels to obtain a single value per 0.5 Hz and then logarithmically scaled. Statistical tests were performed to detect significant differences between groups. Additionally, PSD data was aggregated into specific frequency bands: delta (1-4 Hz), theta (4-8 Hz), alpha (8-13 Hz), beta (13-30 Hz) and gamma (30-40 Hz) and averaged across epochs. This information was used to plot topomaps for each frequency band in each condition, and topomaps for the statistical test between conditions, identifying channels showing significant differences

The complexity of broadband signals was assessed using the Lempel-Ziv lossless compression algorithm (Schartner et al., 2017a; Timmermann et al., 2019; Cavanna et al., 2022; Pallavicini et al., 2021a). This algorithm divides a binary string into non-overlapping, unique substrings. The greater the diversity within the string, the larger the number of substrings. The total count of these substrings is referred to as the Lempel-Ziv complexity (LZc). The initial step involved computing the instantaneous signal envelope for each epoch and channel, using the Hilbert transform. Following this, Z-score normalization and binarization through a median split were performed. The median split ensured an equal proportion of 1s and 0s across all channels, thereby mitigating biases arising from unbalanced sequences. Finally, Lempel-Ziv complexity (LZc) was calculated for the binarized signal of each epoch and channel, and the values were averaged across epochs to obtain a single value per channel.

### Statistical analyses

Questionnaire results were compared between the dosing conditions using Student’s paired t-test, as implemented in Python’s Scipy library (https://scipy.org). We reported uncorrected p-values, and highlighted instances where p-values remained significant after applying the Benjamini-Hochberg false discovery rate (FDR) correction at α=0.05. Corrections were not applied to instances where statistical independence could not be assumed, such as entropy results derived from multiple thresholds. Effect sizes were calculated using Cohen’s d for parametric tests and rank-biserial correlation (RBC) for non-parametric tests, with a detailed summary provided in Table 1 and Supplementary Table S1. Estimating the appropriate sample size was challenging due to the limited prior research on eye-tracking metrics in psychedelic studies. We estimated a sample size of 23 participants based on an expected medium to large effect size (d = 0.6), which is commonly observed in studies of psychedelics like psilocybin, with statistical power set to 0.8 and a significance level of 0.05, in a two-tailed, within-subjects design. To complement frequentist analyses, Bayesian statistics were employed to assess the relative evidence for the null and alternative hypotheses. We computed the Bayes factor (BF10) favouring the alternative hypothesis, using Python’s pingouin library (https://pingouin-stats.org) (Held & Ott, 2018). Non-parametric Wilcoxon signed-rank tests were used to compare eye-tracking measures and VAS scores across dosing conditions.

Figures display boxplots with the interquartile range, where the median is indicated by a central line. Whiskers extend to 1.5 times the interquartile range, and individual data points from each participant are overlaid as scatter points above the boxplots.

## Results

The experimental condition was correctly unblinded by participants in 42 out of 46 measurement days (91%), with this rate being identical for both the high dose and active control conditions. Details on all self-reported scales and questionnaires can be found in the supplementary material (Table S1). As anticipated, the high dose condition produced significant increases across all items of the MEQ30 and 5D-ASC questionnaires. Furthermore, most negative results had BF10 factors below 1/3, supporting the null hypothesis (i.e., no effect of psilocybin).

### Statistical fixation metrics

The total number of fixations (N), average distance between fixations (ds) and average time between fixations (t), were computed for each stimulus and averaged over all stimuli for each subject and dosage condition, as detailed in the Methodology section. To assess differences between dose conditions, we performed non-parametric paired Wilcoxon signed-rank tests (Figure 2b). The analysis revealed a significant effect of dosing, with the metric *ds* exhibiting lower values under the high dose condition.

**Figure 2.**
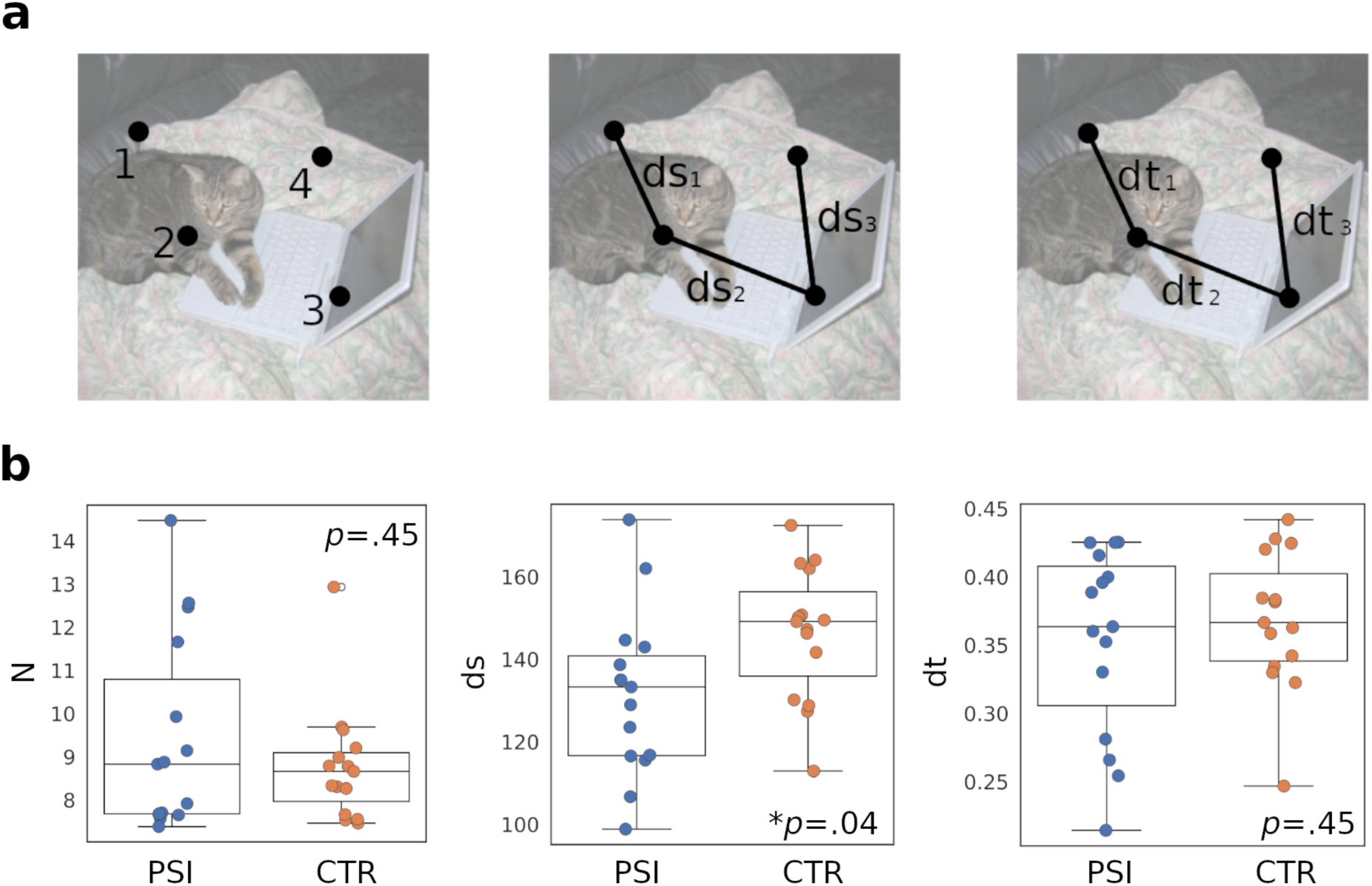
Changes in fixation metrics in high vs. low dose conditions. **a)** Illustration of the statistical fixation metrics that were calculated for each trial. From left to right: number of fixations (N), distance between consecutive fixations (ds) and time between consecutive fixations (dt). **b)** Values of the metrics N, ds, and t, averaged across stimuli for each subject, and compared between the high dose of psilocybin (PSI) vs. the active control (CTR) condition. The p-values (Wilcoxon signed-rank tests) are shown as insets. *p<0.05

### Visual saliency

As described in the Methods section, normative saliency maps were generated using *DeepGaze II* to quantify the visual salience associated with the subjects’ fixations (Figure 3a). The average saliency values across all images were compared between the high-dose psilocybin (PSI) and active control (CTR) conditions. The high-dose condition showed significantly greater saliency compared to the control condition. Furthermore, higher saliency values were also observed in the second, third, and fourth of the five intervals into which the 3-second viewing period was divided, for the high-dose psilocybin (PSI) condition compared to the active control (CTR) condition (Figure 3b).

**Figure 3.**
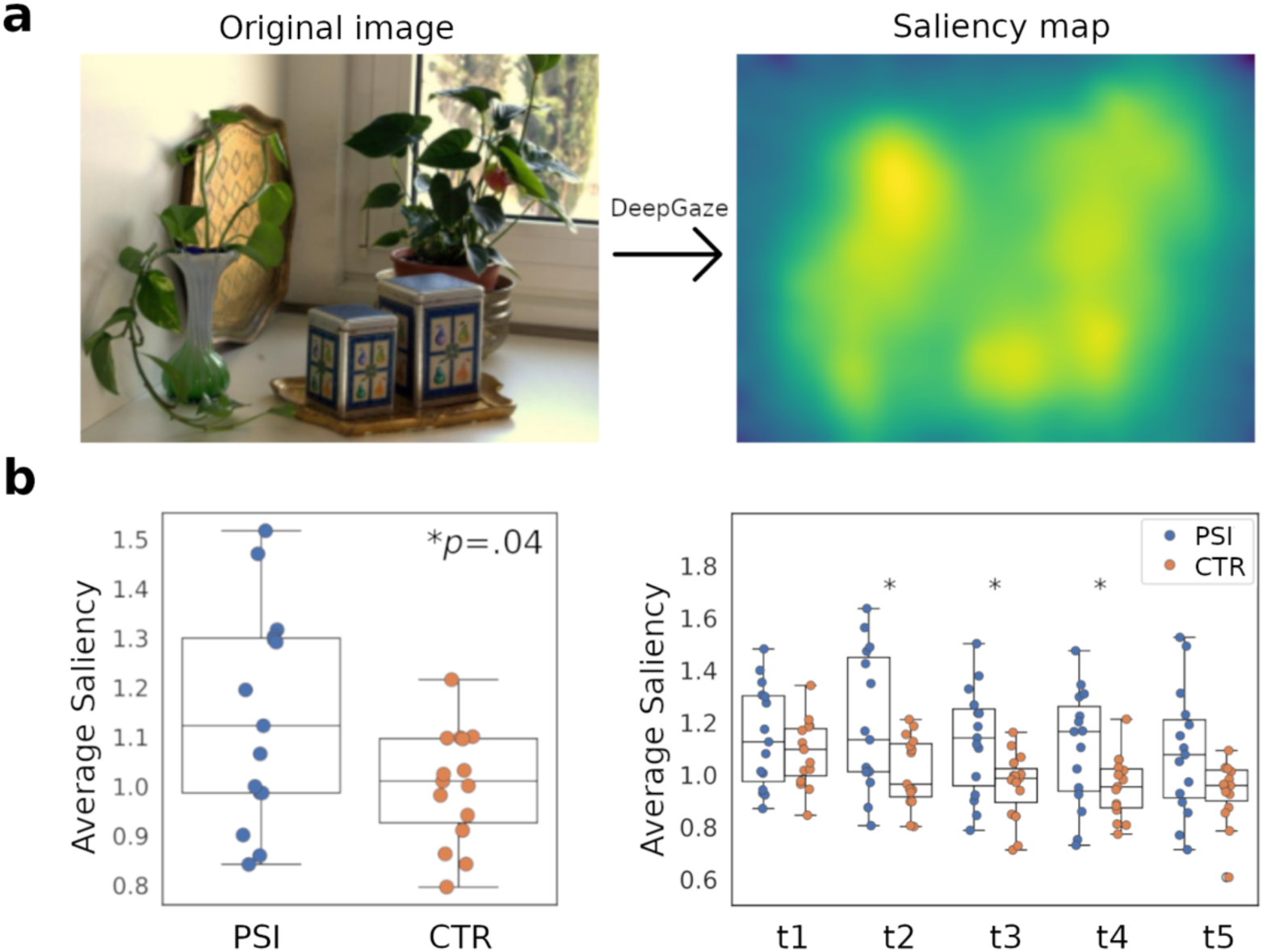
Changes in average saliency under high vs. low dose conditions. **a)** Example of saliency map generation using DeepGaze II. **b)** Average saliency values across stimuli for each subject, compared between the high dose of psilocybin (PSI) and the active control (CTR) condition. Left: overall average across fixations. The p-value (Wilcoxon signed-rank test) is shown as inset. Right: averages calculated for five equal time intervals during the 3-second viewing period. *p<0.05.

### Shannon Entropy in High Salience Regions

The calculation of Shannon Entropy in high salience regions, as described in the Methods section, is illustrated in Figure 4a. The number of clusters detected in the binarized saliency maps varied as a function of the threshold, as shown in the left panel of Figure 4b. To ensure a robust calculation of entropy, we selected a range of thresholds (marked by red lines). Green vertical lines indicate all thresholds within this range where entropy values were significantly different between conditions, with higher entropy consistently observed for the high-dose psilocybin (PSI) condition compared to the active control (CTR) condition. The right panel of Figure 4b illustrates this result for a specific threshold within the selected range.

**Figure 4.**
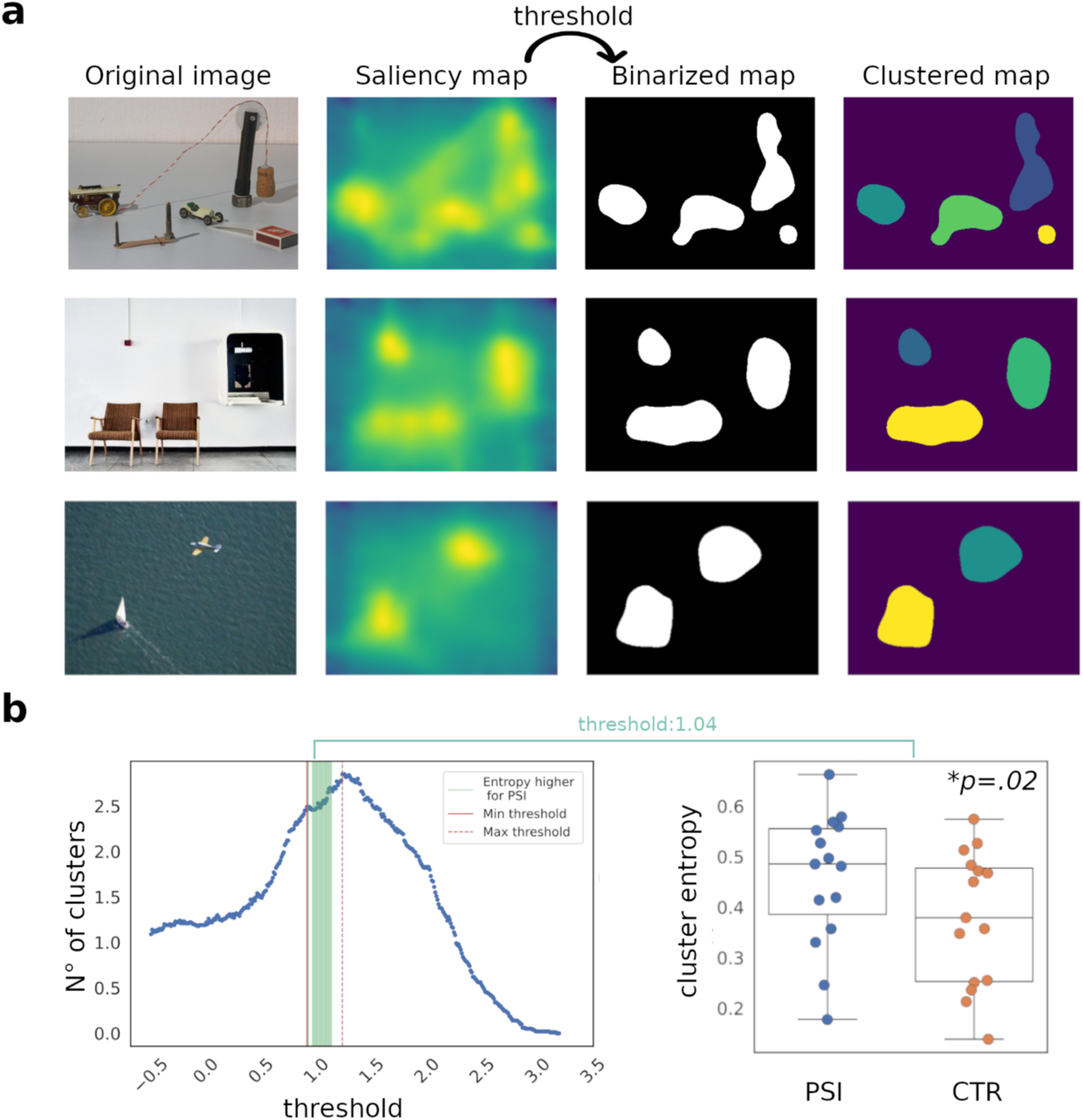
Entropy in high saliency regions. **a)** Examples of high saliency region clusterization: original image, saliency map, binarized map, and clustered map are shown for three different stimuli. **b)** Left: Number of clusters as a function of the threshold. The red lines mark the selected range of thresholds used for entropy calculations, while the green lines indicate thresholds where entropy was significantly different. Right: Example of entropy values for a specific threshold within the selected range, showing significantly higher entropy for the high dose (PSI) condition.

### EEG spectral power and Lempel Ziv complexity

We compared the logarithmic PSD between dose conditions, averaged across all channels, as shown in Figure 5a. Light vertical lines indicate significant differences, while dark vertical lines denote significant differences after multiple comparisons. We observed decreased spectral power for frequencies in the upper part of the delta band (1-5 Hz), and in the alpha (8-12) and beta (12-20) frequency bands. We also observed increased spectral power in the gamma range (>35 Hz) for the high dose condition. Figure 5b displays the topographic distribution of z-scored spectral power for the delta, theta, alpha, and gamma bands, as well as the broadband Lempel-Ziv complexity, for both dose conditions. Additionally, topographic maps of the t-values derived from the statistical tests comparing conditions are also displayed, for the same frequency bands and complexity measure. As shown in the last column of Figure 5b, increased Lempel-Ziv complexity was found in the high dose condition across all channels.

**Figure 5.**
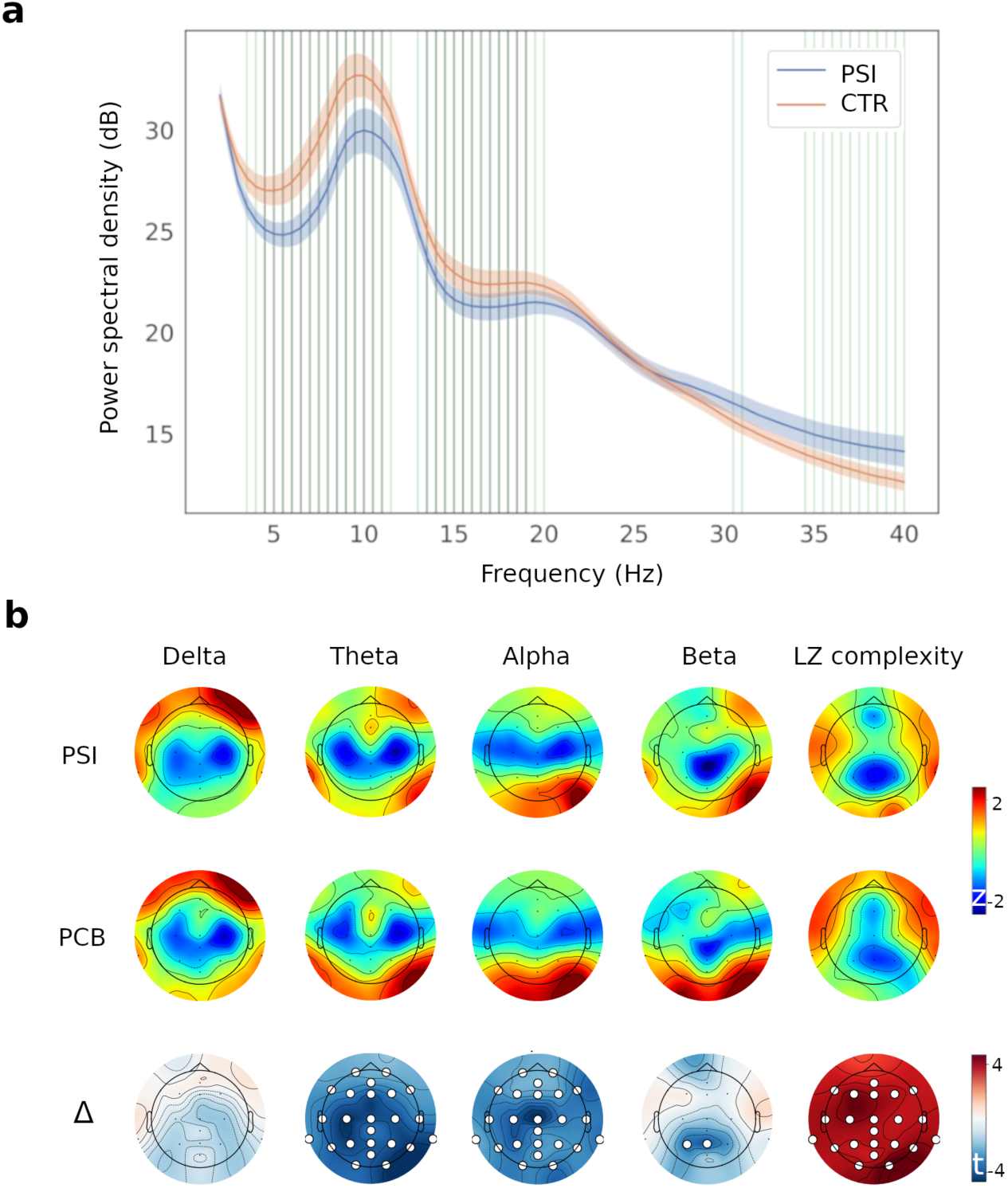
Comparison of spectral power and complexity across dose conditions. (a) Logarithmic PSD averaged across all channels, with light vertical lines indicating significant differences and dark vertical lines highlighting significant differences after multiple comparisons. (b) Topographic maps of spectral power across frequency bands and Lempel-Ziv complexity for both dose conditions, along with topographic maps of t-values from statistical tests comparing conditions.

**Figure 6.**
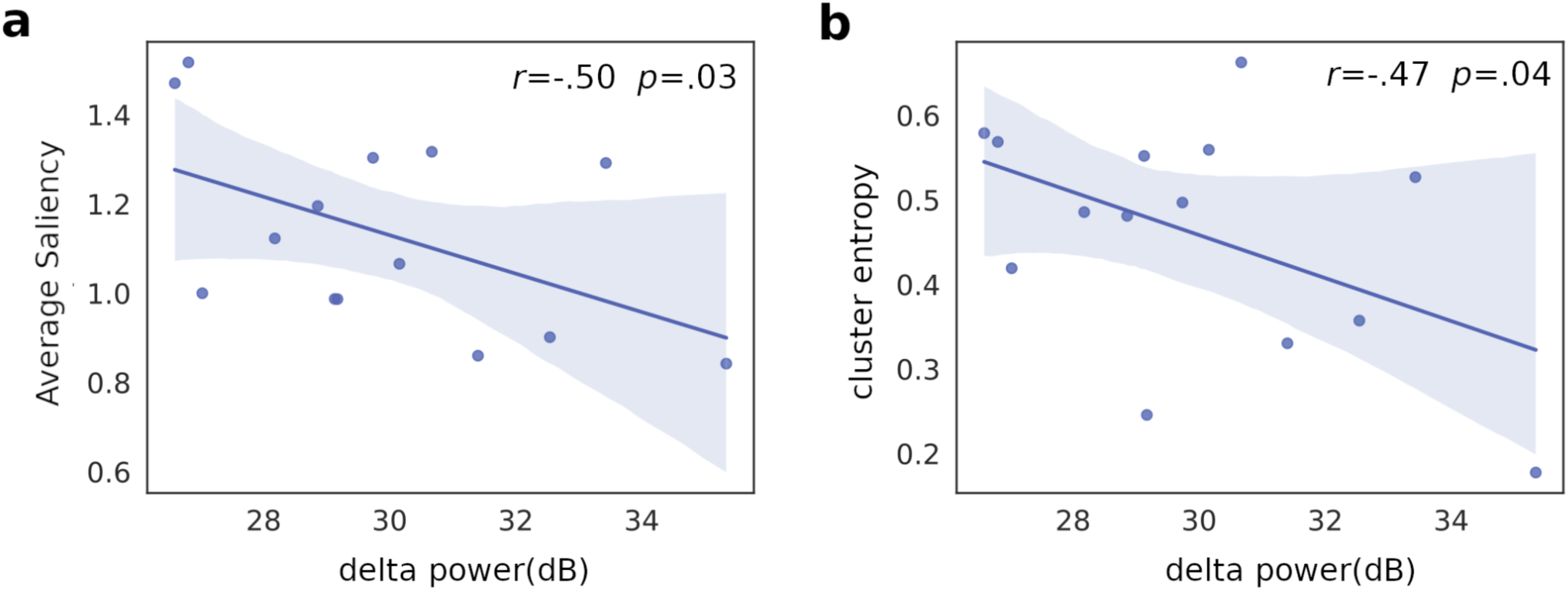
Scatter plots of saliency metrics vs delta power, including the best linear fit in the least squares sense for the high dose condition **a)** Negative correlation between delta power and average saliency **b)** Negative correlation between delta power and high saliency cluster entropy. Shaded areas represent 95% confidence intervals. Pearson’s correlation coefficient (*r*) and corresponding *p*-values are displayed in the insets.

### Correlations between spectral power and behavioral metrics

To investigate the relationship between neural oscillatory activity and visual exploration behavior, we computed correlations between spectral power measures (averaged across all EEG channels) and salience metrics estimated using the *DeepGaze II* model. This analysis revealed significant negative correlations between delta power and two saliency-based metrics: average saliency across all fixations, and the Shannon entropy of the high salience regions explored during visual perception.

## Discussion

This study explores how psilocybin influences initial gaze behavior in natural scenes, focusing on eye-tracking metrics and saliency processing. Our findings indicate that psilocybin induces significant alterations in gaze behavior, characterized by shorter distances between fixations and increased attention toward visually salient regions. These effects are accompanied by higher entropy of the sequence of high saliency regions explored during the visual perception task, suggesting a more dispersed and exploratory scanning strategy under the influence of psilocybin. EEG analyses revealed significant dose-dependent changes in spectral power across frequency bands and Lempel-Ziv complexity. Notably, delta power showed significant negative correlations with both average saliency and cluster entropy in the high dose condition.

The observed reduction in the average distance between fixations aligns with our previous report indicating more localized visual exploration during psychedelic states (Muller et al., 2023). This could reflect a tendency toward hyper-focused attention or difficulty in disengaging from visually compelling stimuli. Additionally, the significant increase in saliency associated with fixations suggests a shift toward more stimulus-driven attention, with salience exerting greater influence over gaze allocation. This change in visual salience processing under the acute effects of psilocybin may reflect a weakening of top-down modulation, allowing bottom-up sensory input to exert a greater influence on perception, a mechanism consistent with the hypothesis of relaxed beliefs under psychedelics (REBUS model) (Carhart-Harris & Friston, 2019).

At the neural level, classical psychedelics are known to disrupt the functional connectivity, cerebral blood flow and oscillatory activity within the Default Mode Network (DMN), a network involved in self-referential thought and internally directed cognition (Gattuso et al., 2023; Carhart-Harris et al., 2012; Muthukumaraswamy et al., 2013; Palhano-Fontes et al., 2015; Lebedev et al., 2015). Psychedelics reduce the modular organization of brain activity by enhancing inter-network connectivity (Roseman et al., 2014; Palhano-Fontes et al., 2015; Tagliazucchi et al., 2014, 2016), decreasing functional segregation and potentially contributing to weakened top-down predictive control, with behavior predominantly guided by stimulus salience.

The salience network, which plays a key role in filtering and prioritizing behaviorally relevant stimuli (Seeley, 2019; Uddin, 2016), is also disrupted under psychedelics, with psilocybin in particular reducing its integrity (Lebedev et al., 2015). The compromised filtering and regulatory functions of the salience network, coupled with the general increase in sensory vividness under psychedelics (Kometer & Vollenweider, 2018), might lead to a greater focus on what naturally stands out in visual perception. This altered attentional state may lead participants to prioritize naturally salient image regions, as perception becomes more influenced by intrinsic stimulus properties rather than learned gaze patterns.

The increase in Shannon entropy within the sequence of high-saliency regions explored during perception further reinforces the idea of less constrained visual exploration under psilocybin. Importantly, this effect is observed specifically within salient regions, rather than across the entire visual field, suggesting that while psilocybin enhances attention toward intrinsically significant stimuli, it also disrupts the conventional, structured way in which these stimuli are typically processed. Under normal conditions, observers exhibit predictable fixation patterns that optimize information gathering based on learned expectations and scene context, prioritizing efficiency and familiarity (Henderson et al., 2007). However, the psilocybin-induced increase in entropy suggests a loosening of these constraints, leading to a more stochastic and dynamically shifting scanning behavior within the most visually dominant regions. This shift aligns with broader findings on psychedelic-induced cognitive flexibility and exploratory behavior (Bălăeţ, 2022). Rather than adhering to habitual attentional routines, individuals under psilocybin engage with high-saliency areas in a less predictable, more fluid manner, potentially allowing for novel interpretations and richer perceptual experiences.

EEG results corroborated the reports of subjective effect intensity, with significant broadband reductions in spectral power, a robust neural marker of psychedelic effects (Muthukumaraswamy et al., 2013; Pallavicini et al., 2021b; Kometer et al., 2013; Riba et al., 2002; Schartner et al., 2017b; Schenberg et al., 2015; Timmermann et al., 2019; Valle et al., 2016; Carhart-Harris et al., 2016). The negative correlations observed between delta power and both average saliency and saliency cluster entropy suggest that neural desynchronization may underlie the increased sensory-driven attentional patterns. Reduced delta power, often associated with diminished top-down control (Harmony, 2013) may therefore reflect a loosening of higher-order constraints on perception, facilitating a shift toward bottom-up sensory dominance.

Our results are also relevant for understanding salience processing in conditions that overlap with the acute effects of psychedelics. The increased attention toward visually salient regions induced by psilocybin aligns with the concept of a ’bottom-up bias’ observed in schizophrenia, where attention is predominantly driven by the physical attributes of stimuli. Recent research indicates that when viewing highly physically salient images, the bottom-up computational model of visual saliency is more accurate in predicting the eye fixations of schizophrenia patients compared to healthy controls (Adámek et al., 2024). Furthermore, individuals with schizophrenia exhibit greater tendency to fixate on regions with high orientation salience, indicating a heightened focus on visual features related to orientation during free-viewing tasks (Yoshida et al., 2024). The observed shift toward salient areas under psilocybin may thus reflect a similar prioritization of bottom-up visual information, paralleling attentional mechanisms seen in schizophrenia. Furthermore, the finding that fixations under psilocybin were more spatially constrained—evidenced by shorter distances between successive fixations—mirrors reports of narrower gaze areas and shorter scanpath lengths in schizophrenia compared to healthy controls (Adámek et al., 2024). This suggests a potential tendency toward localizing attention on salient elements rather than engaging in broader visual exploration.

This study presents distinct advantages stemming from its naturalistic design, favoring ecological validity over controlled laboratory conditions. The self-blinding of the high vs. low doses allowed the subjects to participate in this study as part of their own natural use of the substance, resulting in a balance between the strictly controlled conditions of a standard double-blind study and the uncontrolled nature of a fully naturalistic study (Haijen et al., 2018; Sanz et al., 2022; Tagliazucchi, 2022; van Elk et al., 2022). Previous studies have demonstrated that laboratory settings are associated with increased anxiety and discomfort in subjects under moderate to high doses of serotonergic psychedelics, which may contribute towards decreasing their engagement in the tasks and negatively affect the overall quality of the recorded neural and behavioral data (Studerus et al., 2012). Psychedelics are also known to induce deeply meaningful experiences with long-lasting effects, posing ethical challenges when studying participants in artificial settings under instructions to perform monotonous and repetitive tasks.

Another advantage of our experimental design is the use of visual perception tasks that do not require active participation from the participants, as in tasks where the main outcomes are performance metrics (e.g. accuracies, reaction times). As in our previous study investigating visual behavior during aesthetic perception (Muller et al., 2023), participants were instructed to passively view the stimuli as they would normally do. It has been established that psychedelics compromise different aspects of cognitive function, including focused attention to task instructions, resulting in barriers to experimental paradigms requiring active participation (Bălăeţ, 2022; Bayne & Carter, 2018). While the study of complex behaviors during natural tasks poses significant challenges, especially concerning data analysis, it may contribute to circumventing the limitations associated with tasks that are biased by impaired performance under the acute effects of psychedelics (Tagliazucchi, 2022).

Our choices regarding the experimental protocol also present some limitations that merit careful consideration. The self-blinding protocol lacked controlled sourcing and chemical quantification of the mushroom material. While the psychedelics effects were confirmed by subjective reports and robust objective neural metrics (i.e. broadband reductions of EEG power), variability introduced by lack of control over the effective psilocybin dose cannot be discarded. Another limitation concerns participant unblinding, a persistent methodological challenge in psychedelic research protocols (van Elk et al., 2022), which is manifest even when implementing active control conditions (i.e., a low dose). However, the objective physiological nature of oculomotor metrics provides reasonable assurance that condition awareness minimally influenced the eye movement patterns observed in our study. Another consideration is the relatively small and predominantly male sample, which may limit generalizability. Finally, the exploratory nature of this study has to be taken into account when discussing its limitations. While exploratory investigations serve a crucial function in understudied domains—particularly regarding the sensitivity of ocular metrics to psychedelic-induced behavioral alterations (Jaeger & Halliday, 1998)—they inherently introduce analytical flexibility that may compromise statistical reliability. To mitigate this concern, we restricted our analytical approach to fundamental oculomotor parameters and saliency and entropy measures, guided by established theoretical frameworks including the entropic brain hypothesis and the REBUS model (Carhart-Harris & Friston, 2019). Future research should build upon these preliminary findings through hypothesis-driven, pre-registered research protocols to further elucidate the observed phenomena.

In conclusion, our findings demonstrate that psilocybin significantly alters visual attention and exploration strategies, shifting behavior toward increased focus on salient regions and less predictable gaze patterns. The altered neural and behavioral state induced by psychedelics may favor perception driven more by intrinsic stimulus features than by goal-directed strategies. Notably, the pattern of enhanced sensitivity to salience under psilocybin echoes attentional alterations observed in schizophrenia, suggesting that psychedelics may offer a useful, though indirect, model for studying atypical salience processing.

## Acknowledgements

This work was supported by grants ANID/FONDECYT Regular 1220995 (Chile) and FONDECYT Exploración 13240170.

## Data availability statement

The raw data and stimuli are publicly available on OSF at https://osf.io/5hr7a/?view_only=971855c5c0cd447fbcbece6a9f477631.

## Author contributions

SM and ET designed the experiment, analyzed data, prepared figures and wrote the final version of the manuscript. SM, FC, LAF, NB, TAD and CP conducted the experiments and reviewed the first version of the manuscript.

## Supplementary material

### Self-reported scales and questionnaires

Big Five Inventory (BFI). A validated Spanish version of the inventory assessing five dimensions of personality: neuroticism, extraversion, openness to experience, agreeableness, and conscientiousness (Benet-Martínez & John, 1998). The BFI questionnaire consists of 44 items based on a 5-point Likert scale. Multiple studies suggest that psychedelics are capable of inducing short and long-term changes in personality (Bouso et al., 2018). Moreover, some of these changes (e.g. increased openness) might be contributing factors to the therapeutic effects of psychedelics, as well as to the long-term positive changes in subjective well-being reported by healthy Individuals

Short Suggestibility Scale (SSS). An inventory that assesses suggestibility, created by Kotov (2004), and translated to Spanish by the authors. The questionnaire consists of 21 items based on a 5-point Likert scale. It has been shown that psychedelics can enhance suggestibility in healthy volunteers (Carhart-Harris et al., 2015)

Tellegen Absorption Scale (TAS). A 34-item scale developed to measure the capacity of an individual to become absorbed in the performance of a task (Tellegen & Atkinson, 1974), translated to Spanish by the authors. This psychological construct is positively correlated with the overall intensity of the effects elicited by psychedelic drugs, and also implicated in some of its most intriguing effects, such as the induction of spiritual or mystical experiences(Haijen et al., 2018)

State-Trait Anxiety Inventory (STAI-T / STAI-S). Validated Spanish editions of commonly used scales which measure state anxiety (situational anxiety of a temporary nature) and trait anxiety (stable trait linked to individual characteristics) (Spielberger, 1983). The instrument comprises 40 items and is based on a 4-point Likert scale.

Positive and Negative Affect Schedule (PANAS). A validated Spanish version of a psychometric scale that has been widely used to measure dimensions of affect, both positive and negative (Watson et al., 1988). The instrument consists of 20 affirmations based on a 5-point Likert scale.

Psychological Well-being Scale (BIEPS). A scale used to measure eudaemonic well-being in adults (including dimensions of acceptance, perception of control, social ties, and autonomy and projects) (Castro, 2002) It consists of 13 questions based on a 3-point Likert scale.Originally developed in Spanish.

Pre-ceremony Scale (PRE). Twelve items assessing non-pharmacological contextual factors prior to psychedelic experiences. Using principal component analysis, Haijen et al. showed that the items clustered into three components: set, setting and clear intentions. (Haijen et al., 2018).

Expectation (EXP). Seventeen questions designed to measure expectations of change in the following areas: positive emotions, negative emotions, anxiety, attention, absorption, creativity, perception, problem solving, empathy, memory, energy, sleep, sociability, spirituality, openness, oceanic feeling and substance intake (Cavanna et al., 2022).

Altered States of Consciousness (5D-ASC). Ninety-four items assessing different aspects of altered states of consciousness, understood as temporary deviations from normal waking consciousness. Consists of three main dimensions, each with several lower-order scales. The dimensions and their corresponding scales are: oceanic boundlessness (experience of oneness, spiritual experience, state of bliss, insightfulness), anxious ego dissolution (disembodiment, impaired control and cognition, anxiety), visionary restructuration (complex images, elementary images, audiovisual synesthesia, altered meaning of perceptions) (Studerus et al., 2010).

Mystical Experiences Questionnaire (MEQ-30). Thirty items from which four subscale scores are calculated: mystical, positive mood, transcendence of time and space, and ineffability, which are considered the most relevant and defining aspects of mystical experiences (Barrett et al., 2015).

A visual analogue scale (VAS) was provided to the subjects for the purpose of rating their subjective effects on four occasions throughout the course of the experiment. This scale is an abridged version of a previously employed scale (Cavanna et al., 2022; Pallavicini et al., 2021), which is limited to items pertaining to the effects of the drug on perception. The items were rated on a visual analogue scale (VAS) comprising the following statements: "Sounds influence what I see", "My sense of size and space is distorted", "I feel unusual bodily sensations", "I see geometric patterns", "Edges seem warped", "I see movement in things that aren’t really moving" and "Things look strange". The outcome of this scale is presented in Figure S1.

**Fig S1:**
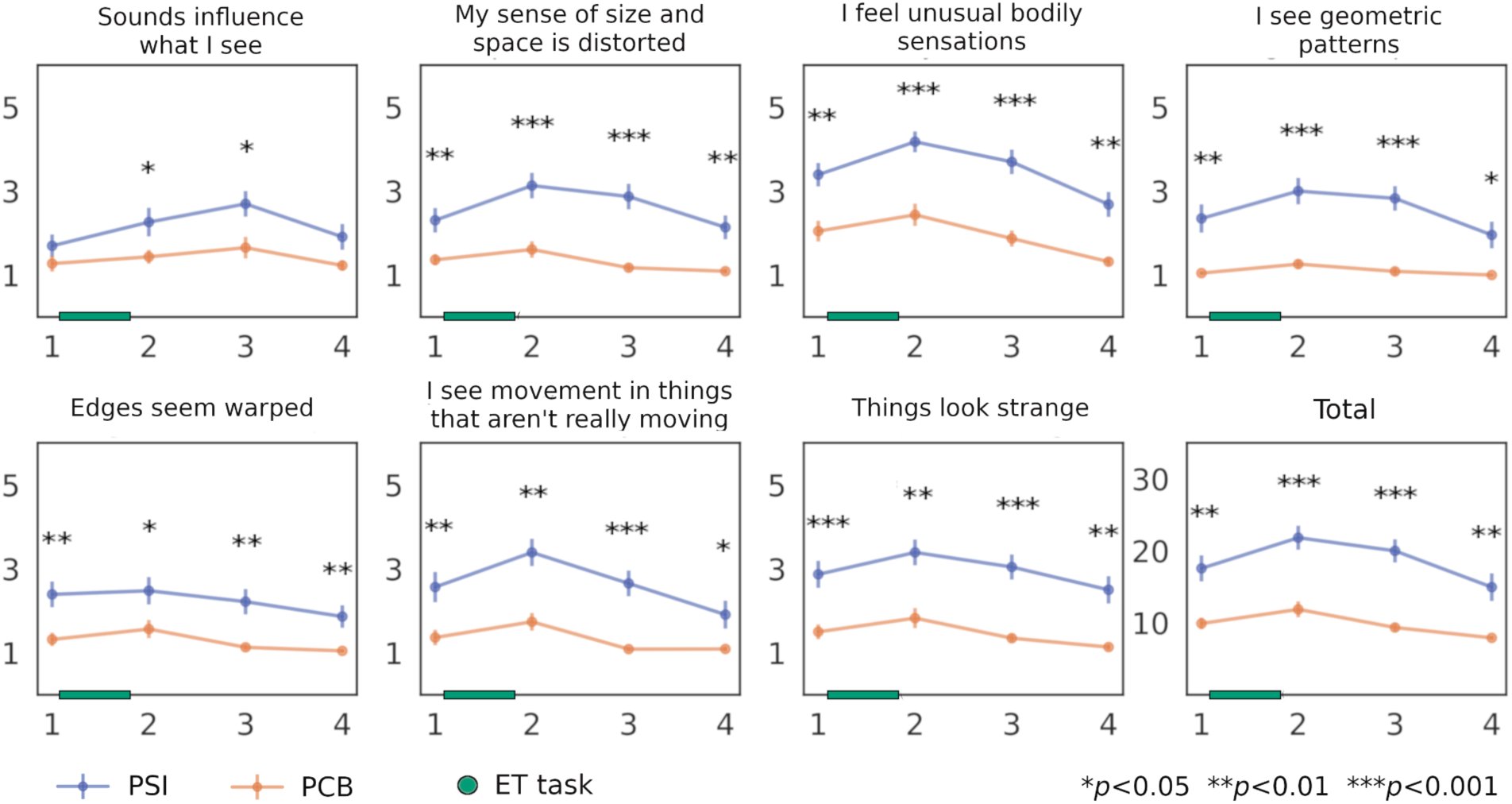
Acute effects measured using individual VAS items and the overall intensity of the experience given by the sum of all the items. Results are shown for each measurement time point, with consecutive measurements separated by one hour. The points indicate the mean across participants and the vertical lines the standard error of the mean (SEM). The time during which the eye tracking task (ET) took place is indicated in the x-axis. Statistical significance is indicated using asterisks (Wilcoxon signed-rank tests). *p<0.05 (Benjamini-Hochberg FDR correction).

### 2. Summary of statistical analyses

The results of all the self-reported scales and questionnaires are presented in Table S1. The results of the acute effects measurements are presented in Table S2.

**Table S1.**
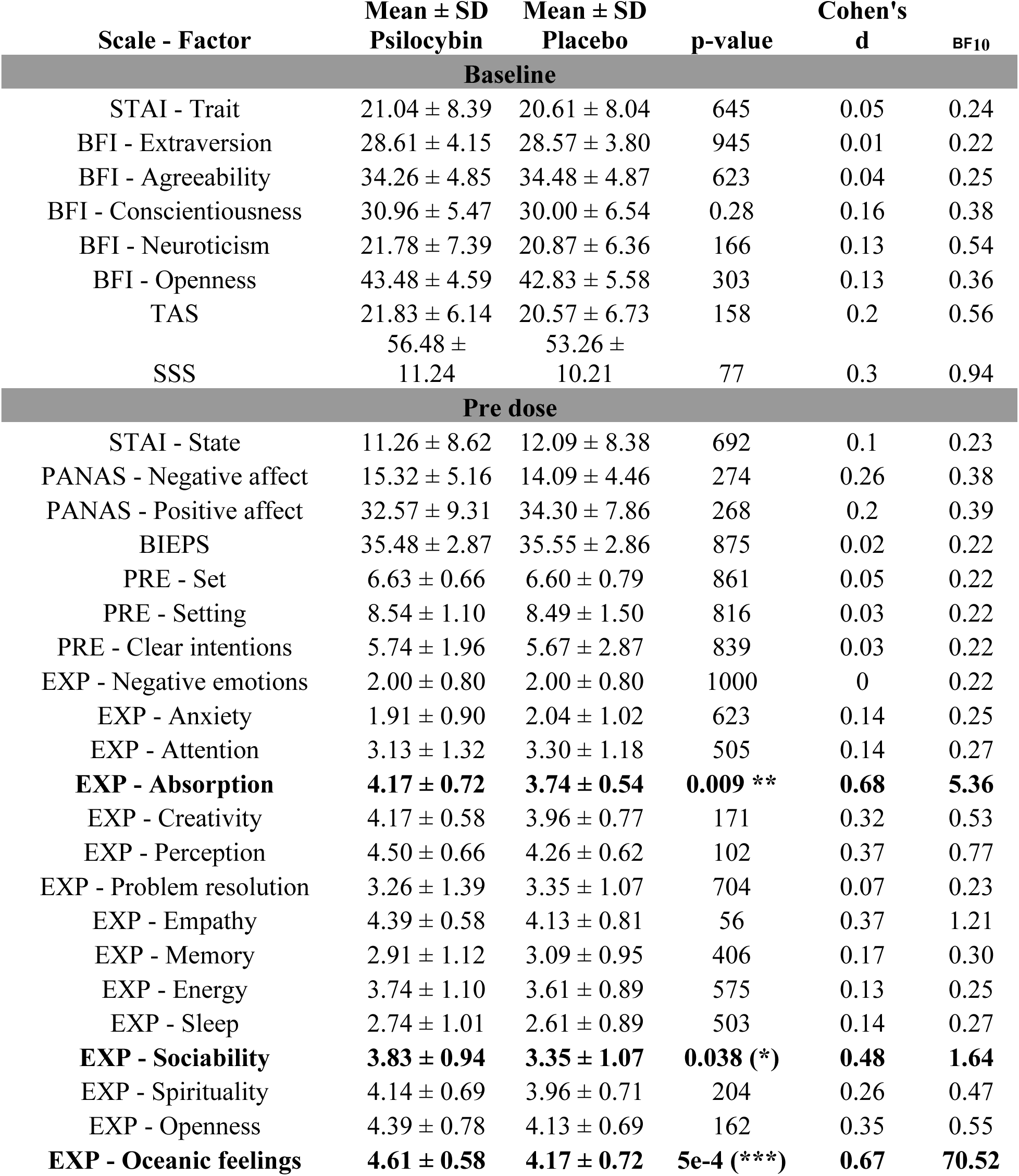
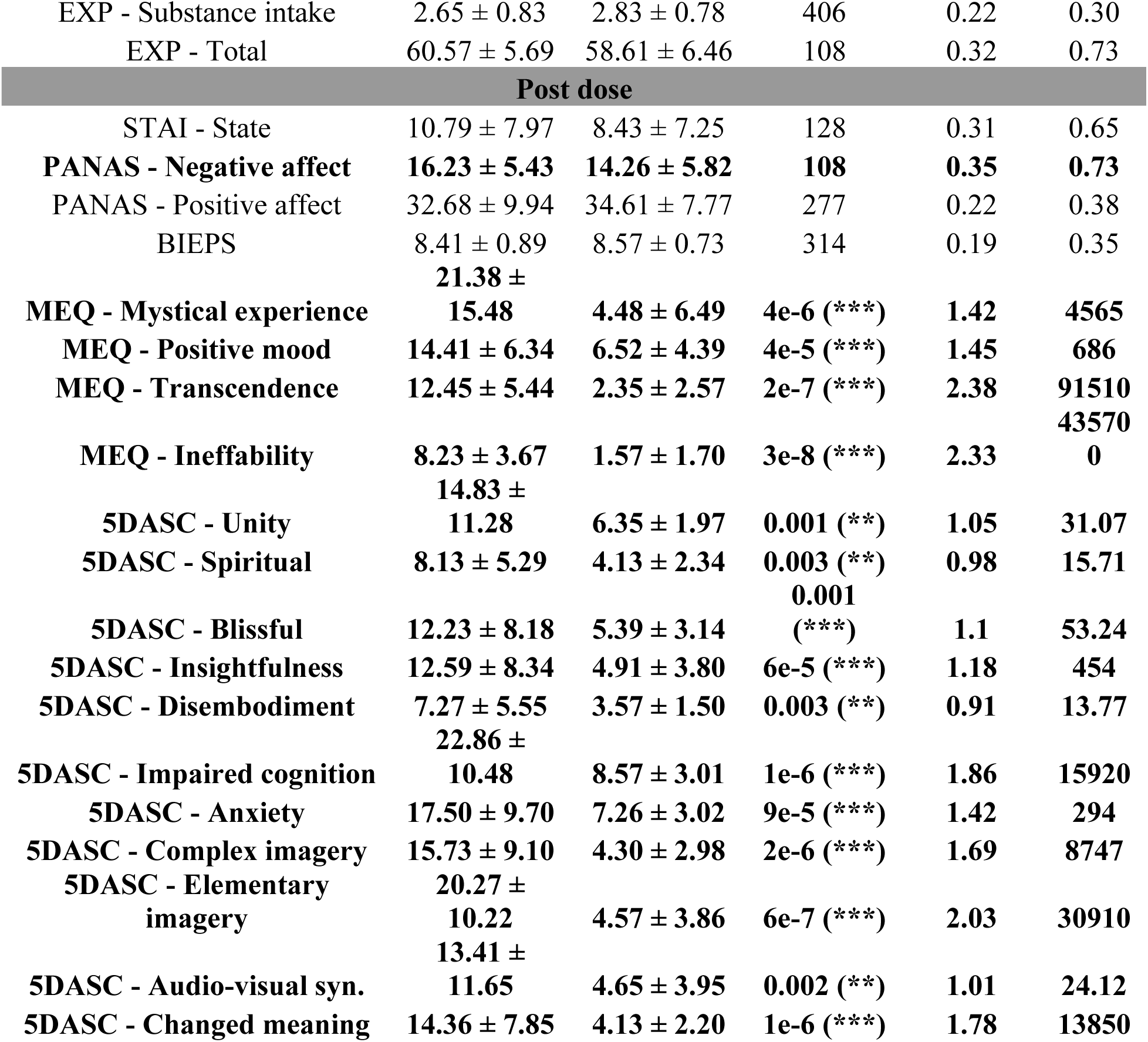
Baseline, Pre dose, Acute effects and Post dose outcomes are presented as mean ± standard deviation. Statistical significance is indicated using asterisks and computed using Student’s t-test.

**Table S2.**
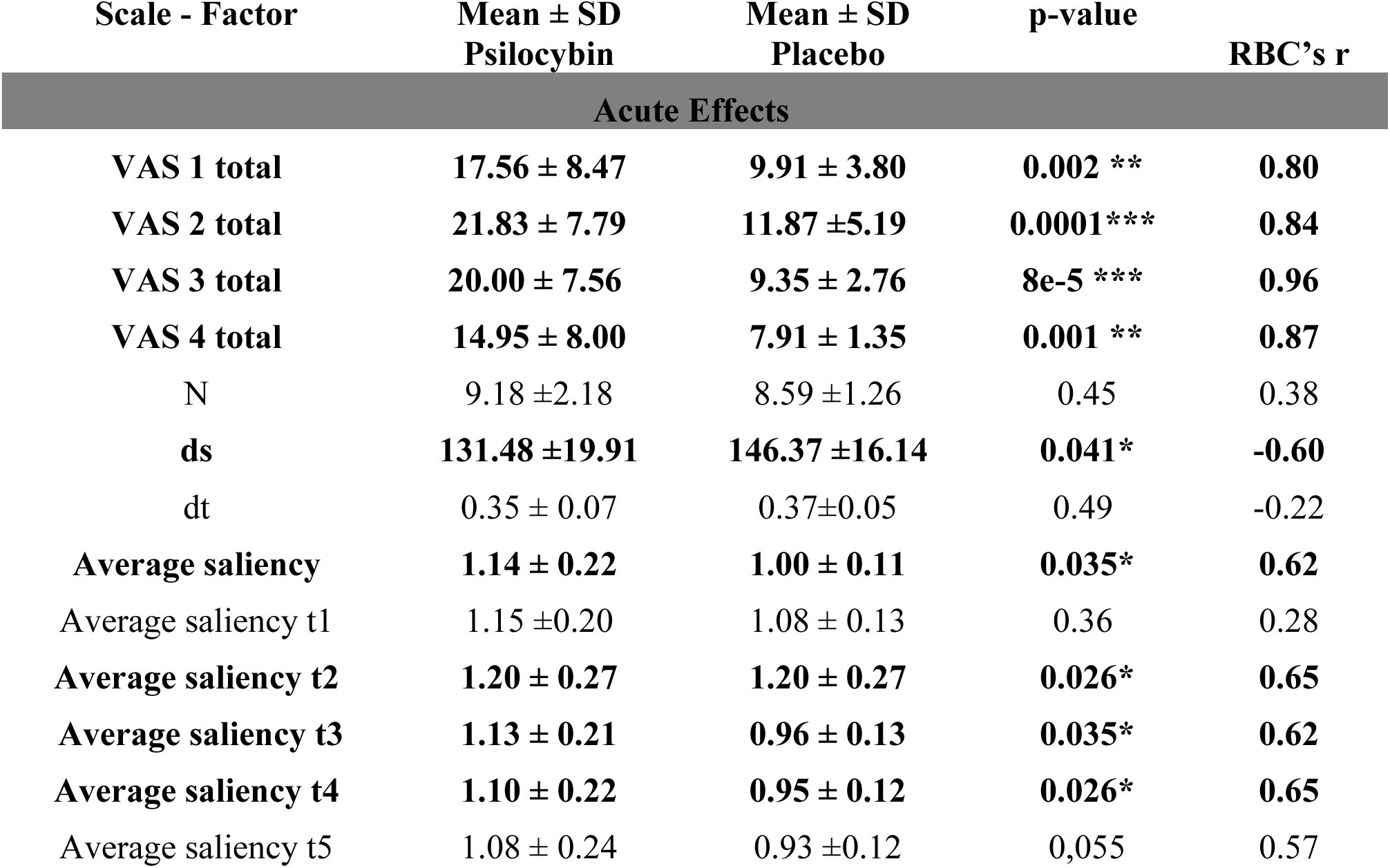
Acute effects outcomes are presented as mean ± standard deviation.

| Scale - Factor | Mean ± SD<br>Psilocybin | Mean ± SD<br>Placebo | p-value | RBC's r |
| --- | --- | --- | --- | --- |
| **Acute Effects** |  |  |  |  |
| **VAS 1 total** | **17.56 ± 8.47** | **9.91 ± 3.80** | **0.002 \*\*** | **0.80** |
| **VAS 2 total** | **21.83 ± 7.79** | **11.87 ± 5.19** | **0.0001\*\*\*** | **0.84** |
| **VAS 3 total** | **20.00 ± 7.56** | **9.35 ± 2.76** | **8e-5 \*\*\*** | **0.96** |
| **VAS 4 total** | **14.95 ± 8.00** | **7.91 ± 1.35** | **0.001 \*\*** | **0.87** |
| N | 9.18 ± 2.18 | 8.59 ± 1.26 | 0.45 | 0.38 |
| **ds** | **131.48 ± 19.91** | **146.37 ± 16.14** | **0.041\*** | **-0.60** |
| dt | 0.35 ± 0.07 | 0.37 ± 0.05 | 0.49 | -0.22 |
| **Average saliency** | **1.14 ± 0.22** | **1.00 ± 0.11** | **0.035\*** | **0.62** |
| Average saliency t1 | 1.15 ± 0.20 | 1.08 ± 0.13 | 0.36 | 0.28 |
| **Average saliency t2** | **1.20 ± 0.27** | **1.20 ± 0.27** | **0.026\*** | **0.65** |
| **Average saliency t3** | **1.13 ± 0.21** | **0.96 ± 0.13** | **0.035\*** | **0.62** |
| **Average saliency t4** | **1.10 ± 0.22** | **0.95 ± 0.12** | **0.026\*** | **0.65** |
| Average saliency t5 | 1.08 ± 0.24 | 0.93 ± 0.12 | 0.055 | 0.57 |

